# Minimizing methane emissions during the degradation of sewage sludge in a sulfate-rich bioreactor

**DOI:** 10.64898/2026.06.23.733557

**Authors:** Gage R. Coon, Oliver Jagoutz, Tanja Bosak

## Abstract

Simultaneous removal of organic waste and industrial gypsum was assessed in continuous flow-through bioreactors that treat sulfate-rich sewage sludge. Metabolic fluxes, the composition of microbial communities, and profiles of organic matter in the presence of different organic loads were tracked over ∼190 days. The addition of a pre-enriched microbial community enhanced the rates of sulfate reduction during the establishment of the sludge blanket, but microbial diversity in established reactors depended primarily on organic loading. Organic removal rates were comparable to those in standard anaerobic digesters, but methane production accounted for ∼1% of electron flow compared to >70% in traditional systems. Stoichiometric analyses revealed that molar COD: sulfate ratios below ∼1 favored complete oxidation of acetate by sulfate-reducing bacteria (SRB) and those above ∼2.1 permitted either complete or incomplete oxidation, allowing sulfate reduction and methanogenesis to co-occur. Sequencing of the 16S rRNA confirmed these trends by revealing that the faster-growing SRB that do not oxidize acetate were more abundant at higher organic loads and during the establishment of the sludge blanket, whereas complete oxidizers became more abundant when the molar COD: sulfate ratio was ≤3.2. In reactors that had been seeded with the pre-enriched communities, acetate-oxidizing SRB became prevalent over the incomplete oxidizers 25-50 days earlier. These results enable targeted design and control of microbial processes and bioreactors that remove waste organics and gypsum while producing less methane due to the competition for acetate between methanogenic archaea and SRB that oxidize acetate.

## 1. Introduction

In modern wastewater treatment, organic matter from sewage sludge is transformed into methane via methanogenesis. Whether this methane is vented directly, flared into CO_2_ (Oshita et al., 2014), or lost as fugitive leaks even in systems designed to recover methane (de Jong et al., 2026), methane remains a substantial contributor to overall greenhouse gas emissions. Adding waste gypsum as a source of sulfate into existing sewage treatment infrastructure can mitigate these emissions. With over 300 million tons produced annually and stacked indefinitely or discharged into the ocean (Bilal et al., 2023), waste gypsum presents environmental challenges associated with disposal issues. This has motivated the search for economically viable strategies to valorize waste gypsum. By adding gypsum directly to anaerobic digesters, electron donors could be redirected from methane production to sulfate reduction (Coon et al., 2026; Murphy et al., 2026). The resulting sulfidic alkaline solutions can be used to recover elemental sulfur (Alsanea et al., 2024; Bounaga et al., 2022; Danouche et al., 2023).

Sulfate-reducing bacteria (SRB) and methanogenic archaea frequently interact in anaerobic environments. The thermodynamic favorability of sulfate reduction predicts the exclusion of methanogens from sulfate-rich environments (Claypool and Kaplan, 1974; LaRowe et al., 2020; LaRowe and Van Cappellen, 2011). However, the Gibbs energy yields of sulfate and methanogenesis can overlap in some environments (LaRowe and Amend, 2014, 2015) and the two metabolisms coexist in marine sediment cores (D’Hondt et al., 2002; Reeburgh, 2007).

Studies of simultaneous sulfate reduction and methanogenesis in laboratory cultures revealed direct competition for H_2_ or acetate (Oremland and Taylor, 1978; Schönheit et al., 1982), and the ability of SRB to outcompete methanogens due to a lower half-saturation constant for hydrogen uptake (Lovley et al., 1982). However, SRB and methanogens may also coexist due to: (1) non-competitive substrate use; (2) direct coupling of intermediates produced by fermentation with methanogenesis; and (3) the flow of SRB-derived organic intermediates such as acetate toward methanogens (Ozuolmez et al., 2015). Non-competitive substrate use occurs when taxonomic groups utilize different, non-overlapping substrates. For example, methanogens can oxidize methyl compounds (Kiene et al., 1986; Lovley and Klug, 1983), and SRB can oxidize propionate (Uberoi and Bhattacharya, 1997). Direct coupling of intermediates is known to occur with acetate produced by fermenters or (homo)acetogens (Dar et al., 2008; Karekar et al., 2022), hydrogen produced by hydrogen-producing fermenters (Oremland and Taylor, 1978), or methyl-compounds produced by methyl-producing fermentation pathways (Bueno de Mesquita et al., 2023; Xiao et al., 2018, 2017; Zhuang et al., 2018, 2016).

The co-existence of SRB and methanogens is reported in areas with high organic loading and excess organic matter and electron donors (Coon et al., 2024, 2023; Egger et al., 2016; Ozuolmez et al., 2015; Sela-Adler et al., 2017). For example, when substrates that are not utilized by methanogens (e.g., lactate) occur at high concentrations, they can stimulate methane production by promoting syntrophic interactions (Sela-Adler et al., 2017). The composition of complex organic sources also controls the composition of fermenting communities capable of degrading such substrates (Hu et al., 2025). Thus, both the organic loading, i.e., the overall availability of reducing equivalents, and the organic composition should shape the competition and cooperation between fermentative microbes, sulfate reducers, and methanogens in bioreactors.

Bioreactors that remove municipal organic waste typically do not contain high concentrations of sulfate, but can be designed to reduce sulfate from seawater while removing municipal (Hao et al., 2013; Lu et al., 2012a, 2012b; Wang et al., 2011, 2009; Wu et al., 2016) or industrial waste (Kijjanapanich et al., 2014; Plyatsuk and Chernish, 2014; Rzeczycka et al., 2010; Zhang and Wang, 2014; Zhang et al., 2022). In such bioreactors, the competition between SRB and methanogens becomes increasingly important because engineered digestion systems contain both SRB and methanogenic archaea (Coon et al., 2026; Murphy et al., 2026; Plyatsuk and Chernysh, 2016) and operate at high organic loadings up a molar COD: sulfate ratio of ∼6 (Wang et al., 2011; Wu et al., 2016). Under these conditions, the coexistence of SRB and methanogens likely depends on non-competitive substrate use (mechanism 1) or the flow of SRB-derived intermediates such as acetate to methanogens (mechanism 3). Recent batch culture experiments that explored the degradation of municipal waste in the presence of waste gypsum identified acetate as the key intermediate (Coon et al., 2026), highlighting the competition for acetate between SRB and methanogens as the likely lever on methane production.

Here, we seek to identify conditions and microbial interactions that minimize methane fluxes while reducing the chemical oxygen demand (COD) in a continuous bioreactor system that processes municipal waste. Tracking metabolic fluxes, microbial community composition, and organic matter profiles as a function of organic loads in three parallel bioreactors over ∼190 days identified optimal ratios of organic loading-to-sulfate that removed COD at rates comparable to those in large-scale methanogenic reactors, but reduced methane emissions. Analyses of microbial community composition showed that lower organic loading led to higher percentage of sequences of sulfate-reducing bacteria that oxidize acetate (CO-SRB) over incomplete oxidizers that excrete acetate (IO-SRB) that is then used by methanogens. The addition of a pre-enriched inoculum to the bioreactors expedited the establishment of CO-SRB and stable conditions relative to the reactors that contained sewage sludge, but lacked a seed community.

## 2. Methods

### 2.1 Bioreactor setup

The upward flow in upflow anaerobic sludge blanket (UASB) bioreactors enables biomass retention that allows settling and accumulation of microbial granules in a stable sludge blanket characterized by high cell densities and metabolic output (Omil et al., 1996). To assess organic removal rates and monitor microbial community composition and activity, three UASB bioreactors (called BR1–3, see Table 2) were designed based on a lactate-fed reactor by Salo et al. (2018), but now feeding the reactor with diluted sewage sludge. The reactors were 0.7 L, 400 mm long, 50 mm wide glass cylinders with jackets for optional temperature control. The reactors were operated at room temperature (20°C–22°C) (Fig. S1). To increase the surface area and stability of the sludge blanket, 350 mL of 4 mm diameter borosilicate glass beads were added to each reactor, yielding a total effective reactor volume of 0.41 L. A marble barely larger than the bottom influent port sat at the bottom to prevent the movement of the 4 mm beads into the influent tubing. A glass airlock (0.175 L volume, 100 mm length, 50 mm inner diameter, 2 mm wall thickness) was added between the reactors and effluent waste to decrease exposure of the reactor to oxygen. All glass sections of the bioreactors were covered by aluminum foil during operation to prevent the growth of photosynthetic microbes (Fig. S1).

Peristaltic pumps pumped and recirculated the medium through oxygen-impermeable tubing (PharMed BPT and Tygon, ¼”). Sewage sludge often clogged the medium inflow tubing (PharMed BPT, 2.8 mm), prompting periodical replacement and manual squeezing along the length of the tubing to dislodge any clumps. Because of this, the flow through the reactor was monitored by measuring the volume of the effluent in the waste container. The flow was stable when the expected volume of effluent was measured in the final container to within ± 10% for the desired hydraulic retention time and the reactor was only sampled when the flow was stable for three or more days. Half-liter Tedlar bags (Restek catalog no. 22049) were installed to collect gases produced by the bioreactors. Fresh medium was stored in a 4°C fridge and stirred by a stir plate. Rubber weights were added to the ends of rubber tubing to keep it submerged in the container and available for medium intake.

### 2.2 Media composition and preparation

Gypsum-saturated water was created by dissolving excess gypsum in tap water, stirring the solution on a stir plate at approximately 400 rpm for an hour, letting excess gypsum settle for an hour, then decanting the saturated fluid into another container. Thickened sewage sludge and final treated effluent were obtained from Deer Island Wastewater Treatment Plant, operated by the Massachusetts Water Resources Authority (MWRA). The COD of the thickened sewage sludge ranged from 74 to 92 g/L, whereas the final treated effluent had a COD of approximately 0.1 g/L.

Different organic loading to sulfate ratios were established by diluting thickened sludge with water in the presence of powdered or dissolved gypsum. Mixture A contained thickened sludge diluted with tap water 55 times, sieved through a 0.35 mm mesh, and combined in equal parts with the gypsum-saturated water to a final 110-fold dilution. Because the dilution required a large volume of liquid, treated effluent was replaced by tap water after the first 20 days, with no observable impact of the change on the performance of reactors. Further mixtures amended dilute sewage sludge with gypsum directly rather than using gypsum-saturated water, and used unsieved sludge because, there was no buildup of settled particulate matter at the bottom of the influent container even in the absence of sieving. Mixture B comprised thickened sludge diluted with tap water 55 times and amended with 2.6 g gypsum per L. Mixture C comprised thickened sludge diluted with tap water 30 times and amended with 2 g gypsum per L. Mixture D comprised thickened sludge diluted with tap water 20 times and amended with 1.47 g gypsum per L.

### 2.3 Inoculation of bioreactors

To compare the inter-reactor variability, bioreactors #1 and #2 (BR1 and BR2) were started and operated at identical conditions in all experiments. These experimental replicates were inoculated with 350 mL of liquid from batch cultures of enriched microbial communities that exhibited high rates of sulfate reduction and COD removal in batch cultures (Murphy et al., 2026). This inoculum reduced sulfate while producing minimal methane at molar COD: sulfate ratios of ∼1.5 (Coon et al., 2026; Murphy et al., 2026).

110 mL of mixture A was pipetted in sterile 160 mL serum vials and inoculated with the enrichment culture (10% v/v), capped by gas-impermeable butyl rubber stoppers, and sealed with aluminum crimps. These cultures were grown at 30°C and 135 r.p.m. in the dark for two weeks. When the concentration of sulfide rose above 3 mM, the cultures were centrifuged at 4,000 x g for 5 minutes and the supernatant removed without disturbing the pellet to remove any metabolic products. The pellet was resuspended in 110 mL of fresh mixture A amended with 2.59 g gypsum/L, and two bioreactors, BR1 and BR2, were inoculated by 350 ml of this suspension each.

The headspaces of the reactors were sparged by nitrogen to reduce the concentration of oxygen, and the inoculum was allowed to grow in the bioreactors at room temperature for two weeks without pumping. The bioreactor tubing was clamped shut to minimize the input of fresh air during this time.

A third bioreactor (BR3) was not inoculated to enable comparing microbial communities that grew in the presence and absence of the inoculum. This bioreactor was amended by 350 ml of DI water before pumping.

### 2.4 Sampling

Reactor performance was monitored by sampling throughout the digestion process. Sludge from the sample port on the recirculating line was sampled into a 15 mL centrifuge tube, homogenized by shaking, and frozen at −80°C for later measurements of chemical oxygen demand (COD), pH, and alkalinity. About 2 mL of this homogenized sample were aerobically filtered through a 0.2 µm filter to quantify sulfate and volatile fatty acids. An aliquot of the filtered sample was mixed with 0.05 M zinc acetate immediately upon filtration to stabilize sulfide for later quantification. Filtered samples were stored at 4°C. Samples for methane measurements were prepared as follows: a 5 mL sample from the sample port was quickly added to a 14 mL serum vial containing 2 mL of 1 M KOH, shaken, capped with a butyl rubber stopper, and sealed with an aluminum crimp seal. Methane was also sampled from the Tedlar gas bag, although there was no observable pressure buildup or volume increase, suggesting negligible degassing from the reactor. Thus, methane concentrations in the sampled sludge should be representative of methane produced therein. An additional 1 mL of the sludge was sampled into sterile microcentrifuge tubes for DNA extraction and stored at −80°C. The same sampling procedure was applied to the fresh medium and the final digested waste effluent. To ensure reliability of the pumping, the residence times of fluids in the reactors were monitored by collecting the waste effluent and measuring its volume over time.

### 2.5 Continuous operation of bioreactors

The continuous operation of the bioreactors began after two weeks of inoculum growth in the absence of pumping and flow. Experiments examined the performance of the microbial communities as a function of different COD: sulfate ratios obtained by mixing sewage sludge, water and gypsum at different ratios (Table 1). Table 2 shows the timing and chemistry of tested conditions.

**Table 1:** Composition of influent mixtures. Values shown are averages of three measurements.

| Condition | COD (g/L) | SO<sub>4</sub><sup>2-</sup> (g/L) | COD: SO<sub>4</sub><sup>2-</sup> (w/w) | COD: SO<sub>4</sub><sup>2-</sup> (mol/mol) | pH | Alkalinity (ppm as CaCO<sub>3</sub>) |
| --- | --- | --- | --- | --- | --- | --- |
| A | 0.77 | 0.84 | 0.92 | 2.76 | 7.47 | 95 |
| B | 1.48 | 1.37 | 1.08 | 3.24 | 7.45 | 148 |
| C | 2.95 | 1.37 | 2.15 | 6.45 | 7.62 | 219 |
| D | 3.67 | 1.00 | 3.67 | 11.0 | 6.39 | 265 |

**Table 2:** Experimental conditions over time.

| Time elapsed (days) | HRT | Mixture | OLR (g/L/d) | SLR (g/L/d) | Notes |
| --- | --- | --- | --- | --- | --- |
| -14 to 0 | Batch conditions |  |  |  | BR1: inoculum settled and darkened<br>BR2: inoculum settled and darkened<br>BR3: no inoculum/no growth |
| 0 to 87 | 12 | A | 1.54 | 1.68 | HRT = 24 hr on days 8–11 during adjustments<br>BR1: day 38: full sludge blanket; partial clog near day 87<br>BR2: day 53: full sludge blanket; day 61: minor backflow causing some drainage (~50 mL)<br>BR3: day 53: full sludge blanket; days 58–60: air pumped into BR3 only |
| 87 to 134 |  | B | 2.96 | 2.74 | BR1: day 91: removed clog<br>BR2 and BR3: no notable issues |
| 134 to 172 |  | C | 5.9 | 2.74 | BR1, BR2, and BR3: no notable issues |
| 172 to 186 |  | D | 7.34 | 2.00 | BR1, BR2, and BR3: no notable issues |

Microbial activity in the bioreactors was monitored continuously and was considered stable only after the reactor had operated at the new condition for at least one week (>14 hydraulic retention times). Data reported for each condition represent the average measurements obtained on at least three distinct sampling days at a constant measured hydraulic retention time (HRT) within 10% of the target flow rate.

Sludge blankets in all reactors were deemed to be fully established when the height of the biomass bed reached the recirculating port (Fig. 1). The sludge blanket was allowed to mature for an additional month with continuous flow after formation in mixture A before formal characterization. Samples were collected during this period at the sampling port to track community assembly.

**Figure 1:**
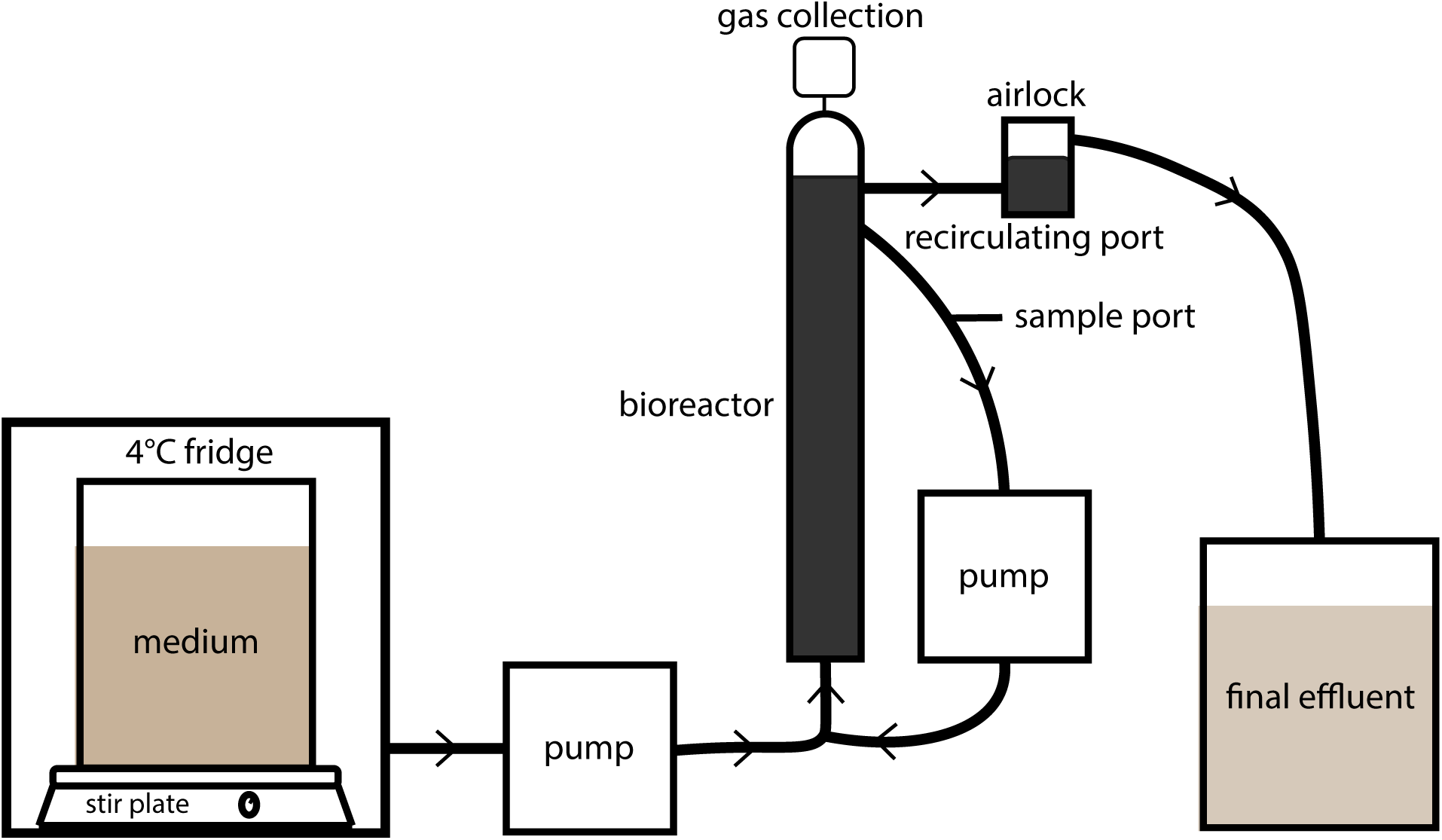
Layout of the bioreactor process design for one of three replicate systems, adapted from Salo et al., 2018.

### 2.6 Sulfide concentrations

Sulfide concentrations were measured by the Cline assay as described here: dx.doi.org/10.17504/protocols.io.6qpvrq1oolmk/v1. Samples were diluted 25-100x in 0.05M zinc acetate and stored at 4°C until measured.

### 2.7 Elemental sulfur concentrations

Samples for measurement of elemental sulfur were taken from precipitates at the top of the bioreactor where the sludge blanket and reactor headspace meet and from precipitates in the airlocks where the fluid and headspace meet. Elemental sulfur concentrations were measured by extracting these 10-25 mg of precipitates and suspending them in 1 mL of water. 0.1 mL of this sample was further suspended into 0.7 mL of 100% acetone and 0.1 mL of 1% w/v zinc acetate. This suspension was incubated for ∼ 10 minutes at 80°C, filtered through a 0.2 µm mesh, and an aliquot of 0.25 mL was dissolved in 0.75 mL of NaCN (1 g NaCN/L dissolved in an acetone: water mix of 19:1 by volume). An aliquot of this solution was mixed with equal parts FeCl_3_ solution (5 g FeCl_3_×6H_2_O/L dissolved in an acetone: water mix of 19:1 by volume). The absorbance of this solution was measured at 465 nm by a spectrophotometer. This method can also detect polysulfides, so the reported concentrations include both elemental sulfur and polysulfides.

### 2.8 Methane concentrations

Methane concentrations were measured using a Shimadzu GC-2014 gas chromatograph with a PDHID and helium carrier gas. 5 mL of sample was quickly placed in a 14 mL serum vial with 2 mL of 1M KOH, then capped and mixed. Dissolved methane ([CH_4aq_], in mM) was calculated using a mass-balance equation to account for both the headspace and residual aqueous methane at equilibrium:

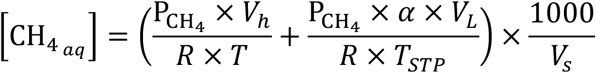

where P_CH4_ is the partial pressure of methane (atm, converted from ppm assuming 1 atm total pressure), V_h_ is the headspace volume (L), R is the universal gas constant (L×atm/mol×K), T is the temperature (K), α is the Bunsen solubility coefficient for methane at the experimental temperature (α = 0.029; Yamamoto et al., 1976), T_STP_ is standard temperature (273.15 K), V_L_ is the total volume of the liquid mixture in the vial (0.007 L), V_s_ is the volume of the original sample (0.005 L), and 1000 is the conversion factor to mM.

### 2.9 Total alkalinity and pH

Alkalinity was measured using a ThermoScientific Orion Total Alkalinity Test Kit (Cat. No. 700010TS). The pH was measured using Vernier Tris-compatible flat pH sensors and AtlasScientific EZO pH sensors.

### 2.10 Chemical oxygen demand (COD)

COD was assessed using a spectrophotometer and a CHEMetrics kit (LR COD Vials 0-1500 ppm, K-7350S, Midland, USA). The COD removal (%) was calculated as the difference in initial and final COD normalized to the initial COD.

### 2.11 Ion concentrations

Sulfate, volatile fatty acids (VFAs), and other ions were measured simultaneously using a dual-channel Shimadzu ion chromatograph equipped with a Shodex SI-52E column. Due to the complexity of sewage, the elution times of many compounds overlapped. To maximize the separation of compounds that are commonly found in solution such as acetate, propionate, lactate, and sulfate, the method employed variable ratios of two influents: A) 5.4 mM sodium carbonate, sparged with N_2_ for 20 minutes, and B) Milli-Q water. The column temperature was set to 50°C for the entire 70-minute run. The flow rate was established at 0.35 mL/min of 30% influent A and 70% influent B for the first 25 minutes, increased to 0.6 mL/min of 100% influent A over the next 15 minutes, increased again to 0.8 mL/min of 100% influent A over the next 10 minutes, held at those conditions for 5 minutes, then returned to the starting conditions (0.35 mL/min, 30% influent A/70% influent B) for 15 minutes. This method was used to determine the concentrations of many fatty acids and other metabolic intermediates such as oxalate, or inorganic compounds such as oxyhalides. Due to the long run time of this method, sulfate was instead quantified using the dual-channel Shimadzu ion chromatograph with the Shodex “Anion Standards (19) Rapid Analysis (SI-90 4E)” method, with the column temperature set to 40°C, for a total run time of 15 minutes.

### 2.12 16S rRNA gene sequencing and analysis

DNA was extracted using the DNeasy PowerSoil Pro Kit (Qiagen, Hilden, Germany) and quantified with Qubit dsDNA HS Assay Kit and Qubit 3 fluorometer (Thermo Fisher Scientific, Waltham, MA, USA). Sequencing was performed on an Illumina MiSeq sequencer using universal primers U515F that target the V4 region of the 16S rRNA gene (5’-GTGCCAGCMGCCGCGGTAA-3’) and E786R (5’-GGACTACHVGGGTWTCTAAT-3’). Sequences were processed with the DADA2 pipeline, and taxonomy was assigned using the MiDAS reference database version 5.3.

## 3. Results

### 3.1 Inoculation and establishment of sludge blanket layers

To determine the time required to establish stable microbial communities that simultaneously remove organic waste and sulfate, we monitored the three UASB reactors after two weeks of batch incubation (see Methods) and after the start of continuous flow of dilute sludge through the system (Day 0). Mixture A was pumped to all bioreactors at a molar COD: sulfate of 2.8 (Table 1); this medium did not contain detectable methane or sulfide before passing through the bioreactors. Sludge blanket layers accumulated for 87 days due to the continuous flow of dilute sludge through each bioreactor (Figure S2).

We hypothesized that providing an inoculum enriched for sulfate reduction would accelerate the onset of sulfate reduction in the bioreactors by reducing the time required for the enrichment and growth of rare sulfate reducers from sewage sludge. As expected, the rates of sulfate reduction increased more rapidly in the inoculated reactors than the uninoculated control reactor after the onset of pumping (Fig. S3). By the time the sludge blankets were fully established, sulfate reduction rates in both inoculated and uninoculated reactors were stable for at least two weeks, though those in the uninoculated reactor remained marginally slower (Fig. S3). These differences can be attributed to the higher initial density of active sulfate-reducing microbes. Methane production remained negligible (< 50 µM/day) across all reactors during this entire phase.

### 3.2 Performance of the bioreactors with established sludge blankets

Following the initial setup and onset of pumping, we characterized the performance of bioreactors as a function of different sulfate and organic loads by monitoring effluent COD, sulfate reduction rates (Fig. 2), and methanogenesis rates (Table S1). Conditions in any reactor were considered as stable only after at least one week of operation (>14 hydraulic retention times). All reported data for any given condition and reactor represent the average measurements obtained on at least three distinct sampling days at a constant measured hydraulic retention time (HRT) within 10% of the target flow rate.

**Figure 2:**
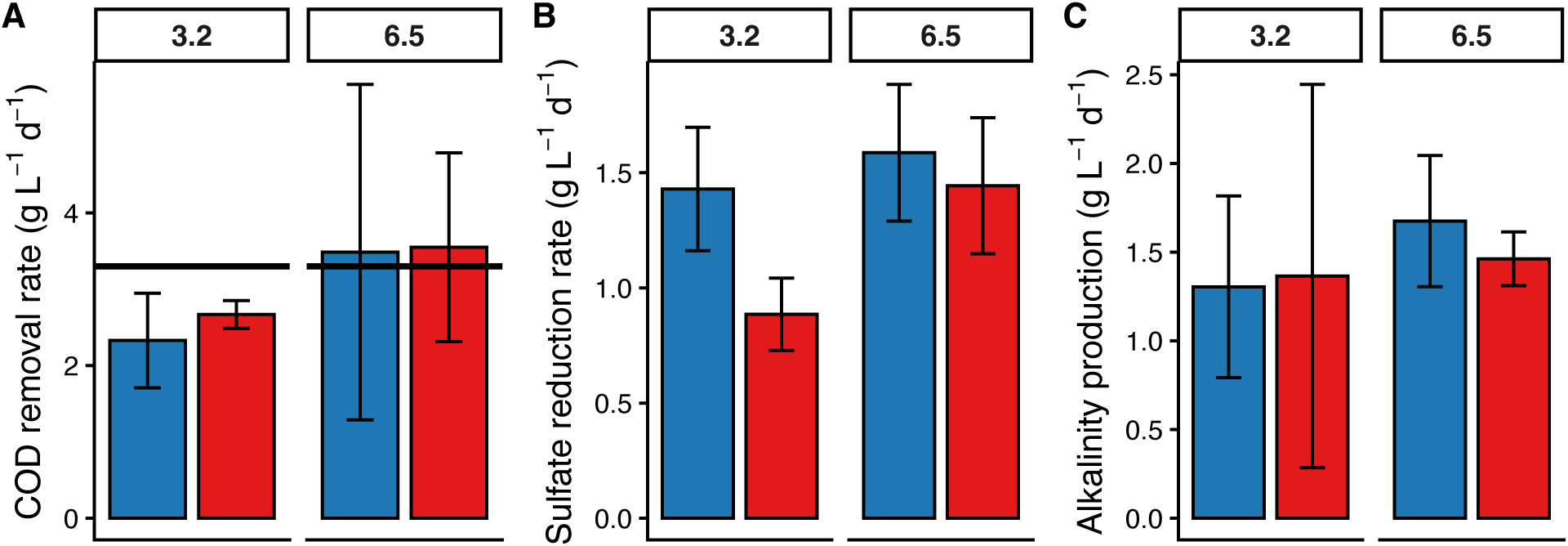
Bioreactor performance after the establishment of the sludge blanket. A) COD removal rate (g/L/d), B) sulfate reduction rate (g/L/d), C) net alkalinity production (g/L/d). Facet labels (3.2 and 6.5) show the tested molar COD: sulfate ratios. Data from duplicate inoculated reactors were averaged (n ≥ 4); temporal replicates from the single uninoculated reactor were averaged instead (n ≥ 2). Error bars show standard deviation. Blue bars show inoculated reactors; red bars show uninoculated controls. The black line on panel A shows the average COD removal rate during sludge digestion at Deer Island Wastewater Treatment Plant, MA, USA.

Once stable sludge blankets were established in all reactors, we observed comparable rates of sulfate reduction and organic matter removal between inoculated and uninoculated reactors across all conditions (Fig. 2). Methane production varied slightly in different conditions but remained low (Table S1).

Rates of COD removal ranged from 1.5 to 4.0 g/L/day, comparable to standard sulfate-free anaerobic digestion systems (Musa et al., 2018). At molar COD: sulfate ratios between 3.2 and 6.5, sulfate remained in excess and reactors removed most of the available oxidizable carbon (Fig. 2A). Conversely, at a COD: sulfate ratio of 11, the effluent did not contain any sulfate, the effluent contained more COD (Table S1), and organic removal rates decreased (Table S1). Organic removal was not characterized at COD: sulfate ratio of 2.8 because the sludge blanket was being established. Sulfate reduction rates were similar at all organic loadings (Fig. 2B, Table S1) and the alkalinity by similar amounts in all reactors when sulfate was not limiting (Fig. 2C).

Methane production remained minimal in all conditions (Table S1). Under standard operating conditions (i.e., excluding sulfate-limiting controls), methanogenesis accounted for ∼1% of the total electron flow (ranging from 0.59% to 1.55%) as COD removal, compared to >70% typically observed in traditional anaerobic digestion. This confirmed successful diversion of electron equivalents toward sulfate reduction rather than methanogenesis.

Acetate is the main electron donor that fuels sulfate reduction and methanogenesis in the batch cultures grown on sewage sludge and sulfate, where it mediates the competition between sulfate reduction and methanogenesis at molar COD: sulfate ratio 4.4 (Coon et al., 2026). To assess the role of this organic acid in the flow-through bioreactors, we monitored concentrations of acetate and other volatile fatty acids at the organic loadings with COD: sulfate molar ratios at 6.5 and 11 to compare sulfate-replete and sulfate-limited conditions (Fig. 3). The final effluent contained more acetate when the COD: sulfate ratio was 11 and the reactors were limited by sulfate. Net production rates, the difference between gross production and acetate oxidation rates, ranged between ∼2-5 mM per day, and the bioreactors produced less than 1 mM methane per day (Fig. 3, Table S1).

**Figure 3:**
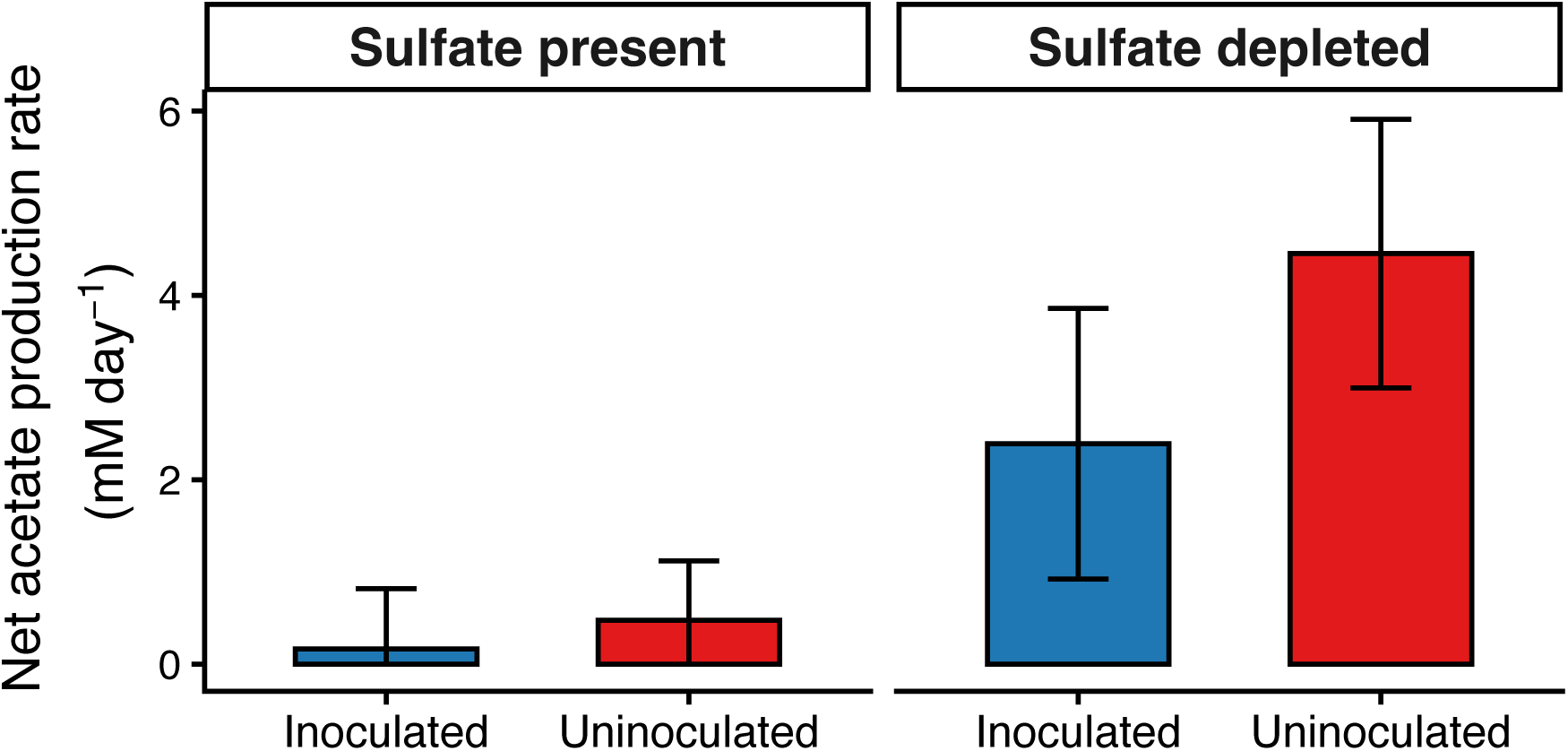
Net acetate production rates in bioreactors limited by organic electron donors (molar COD: sulfate of 6.5) or sulfate (molar COD: sulfate of 11). Data from duplicate inoculated reactors were averaged (n ≥ 6); temporal replicates from the single uninoculated reactor were averaged instead (n ≥ 3). Error bars show the standard deviation.

### 3.3 Microbial community succession

To monitor the compositions of microbial communities and relate the reactor performances to the changes in these communities, we sequenced 16S rRNA genes from all bioreactors at different time points during the operation. Communities established in the reactors were different from those in the influent sewage sludge. The orders most abundant in the influent sludge were *Bacteroidales*, *Lactobacillales*, *Campylobacterales*, *Pseudomonadales*, and *Enterobacterales* (Murphy et al., 2026). Following the onset of pumping and the establishment of the sludge blanket, the reactor communities exhibited similar compositions to one another and contained fermentative *Bacteroidales,* microaerophilic *Campylobacterales*, and the metabolically versatile *Pseudomonadales* (Fig. 4). The retention of *Campylobacterales*, along with aerobic or facultatively anaerobic degraders of recalcitrant organics such as *Burkholderiales*, *Sphingomonadales*, *Pseudomonadales*, and *Flavobacteriales*, suggested that these taxa consumed the dissolved oxygen in the influent sludge and maintained anaerobic conditions to enable the growth of the strictly anaerobic sulfate reducers and methanogens.

**Figure 4:**
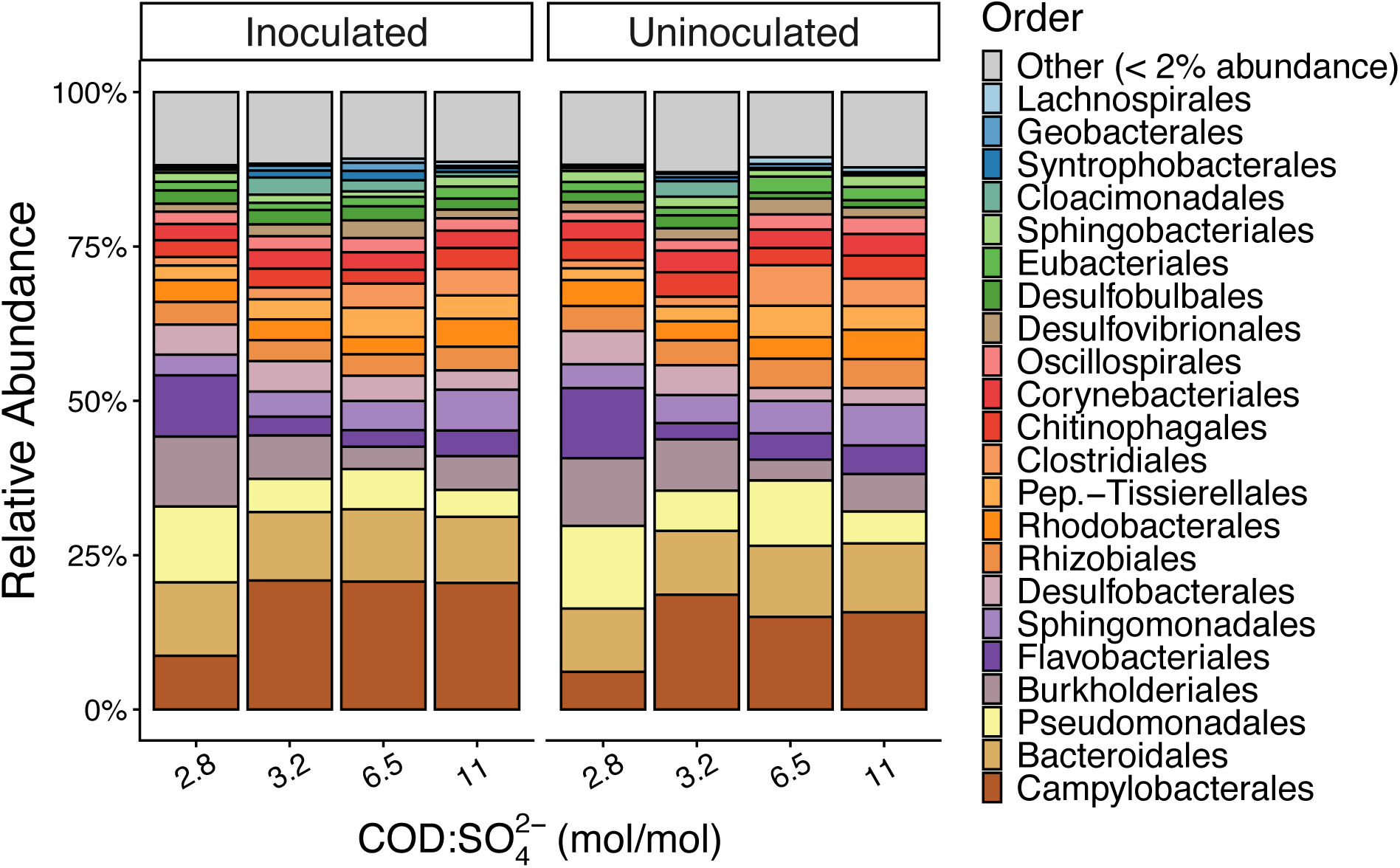
Composition of the microbial communities in bioreactors at the order level. Data show the averages of the final three sampling points for each condition.

### 3.4 Effect of inoculation before the onset of the flow

To understand whether and how the addition of a pre-enriched inoculum altered the trajectory of community assembly, we tracked the beta diversity using non-metric multidimensional scaling (NMDS) analyses of the communities during the first 87 days of pumping under constant conditions (molar COD: sulfate = 2.8; Fig. 5A). At the onset of flow and before the establishment of the sludge blankets, the community structures of the inoculated and uninoculated reactors were similar. The communities in all reactors ultimately converged to a similar community after 70 days (Fig. 5A). Beta diversity remained similar after this time, and changed with the change in the COD: sulfate ratio (Fig. 5B). This indicates that the medium chemistry shapes the community composition more strongly than the presence of a specific starting inoculum.

**Figure 5:**
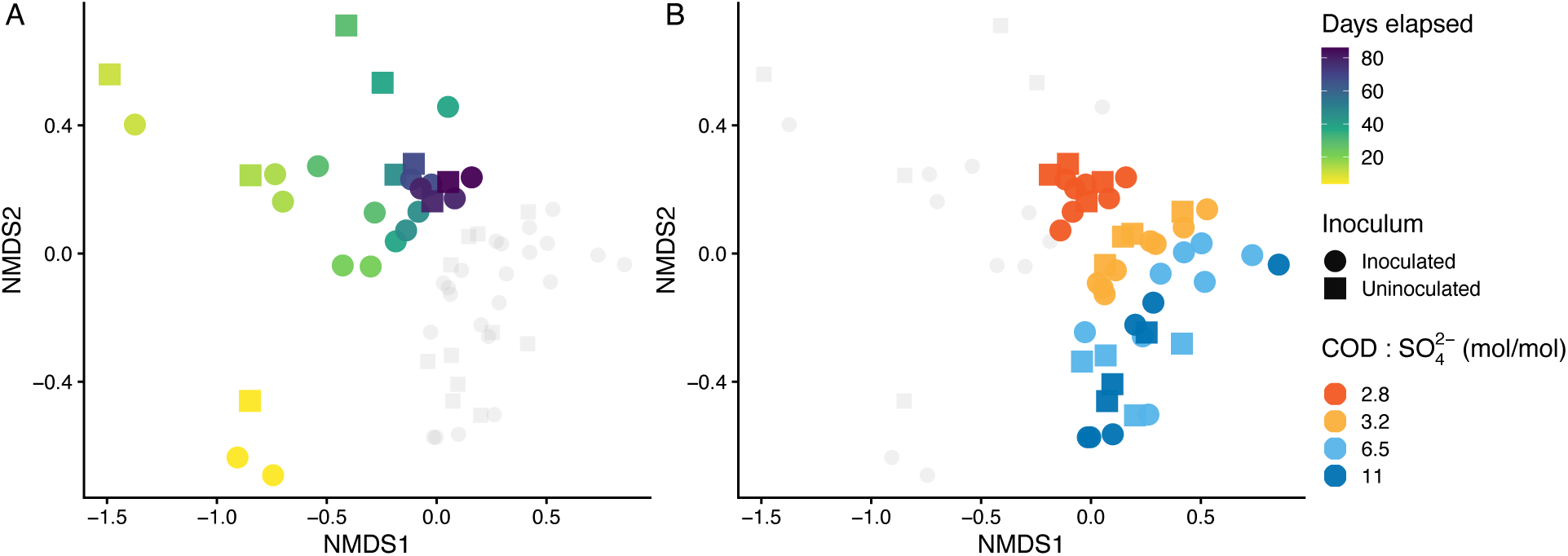
Non-metric multidimensional scaling (NMDS) analysis of microbial community beta diversity. A) Diversity 0–87 days after onset of pumping at molar COD: sulfate = 2.8. B) Comparison of community diversity in the presence of different organic loads. The NMDS ordination was performed on the combined dataset. Colored symbols in both panels highlight specific sample subsets, while light gray symbols show all data points from other sampling times or chemical conditions.

### 3.5 Responses of the microbial community to organic loading

To assess the impact of organic loading on the community structure, we sampled and compared the communities from the reactors that exhibited stable performances at different COD: sulfate ratios (Fig. 5B). The overall community structure at the order level remained relatively stable across the tested organic loads (Fig. 4), particularly when sulfate was not limiting at molar COD: sulfate of 3.2 and 6.5. However, specific taxa responded to the changing organic loads. For example, the relative abundances of *Desulfocapsaceae* decreased at molar COD: sulfate of 6.5 and 11 (Fig. 6A). The relative abundances of aerobic taxa that remove oxygen were higher at molar COD: sulfate of 2.8 (Fig. 4), perhaps in response to the lower organic loading and less reducing conditions. Again, organic loading exerted a stronger influence on community function (Fig. 2) and composition (Fig. 5B) than the addition of the initial inoculum.

**Figure 6:**
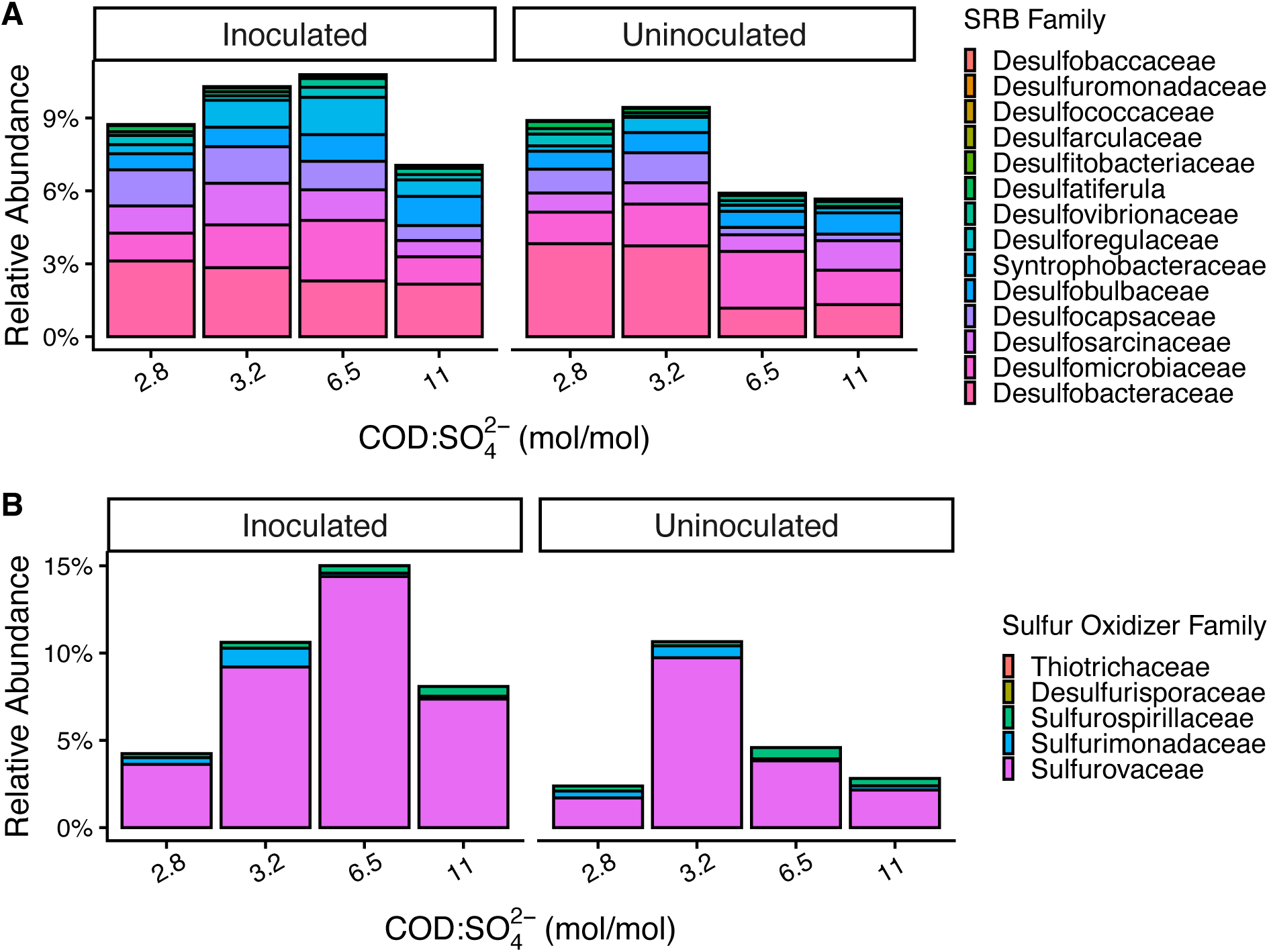
Relative abundances of sulfur-cycling taxa at the family level. A) Sulfate-reducing bacteria (SRB). B) Sulfur-oxidizing bacteria. Data represent the average compositions at the final three sampling times for each condition.

### 3.6 Sulfate-reducing, sulfur-oxidizing, and methanogenic populations

Sulfate-reducing bacteria (SRB) accounted for 5–10% of the total 16S rRNA gene abundances. The SRB community was diverse and multiple families exhibited similar abundances (Fig. 6A). The most abundant taxa were the complete-oxidizing SRB (CO-SRB) *Desulfobacteraceae, Desulfomicrobiaceae* and *Desulfosarcinaceae*. These microbes made up approximately half of the total SRB abundance (Fig. 6A). Incomplete-oxidizing SRB (IO-SRB) were present across all conditions.

The communities represented by the 16S rRNA sequences also contained 5-15% sulfur-oxidizing bacteria, with *Sulfurovaceae* as the most prevalent family (Fig. 6B). High abundances of sulfur oxidizing bacteria coincided with the formation of yellowish-white precipitates at the interface of the sludge and reactor headspace. *Sulfurovaceae* sequences were not detected in incoming sewage sludge and were enriched at the interfaces between anoxic fluids and the oxic headspaces. The precipitates in the reactors contained 4.7% elemental sulfur by mass, those in the airlock contained 1.3% elemental sulfur by mass. The presence of sulfur-oxidizing bacteria and the accumulation of elemental sulfur in areas where the reactor was in contact with air may explain the regularly observed disparity between the amounts of reduced sulfate and the produced sulfide. We did not attempt to address this disparity further so as not to disrupt the sludge blankets and microbial communities in the reactors.

Methane production rates were low in all experimental conditions (Table S1), in keeping with the low relative abundances of methanogenic archaea (< 0.4%, Fig. S4). The detected methanogens were primarily hydrogenotrophic *Methanobacteriaceae* and *Methanoregulaceae*, with a minor population of hydrogen-dependent methylotrophic *Methanomassiliicoccaceae* (Fig. S4).

### 3.7 Abundances of CO-SRB and IO-SRB

We hypothesized that the inoculation of a community enriched in CO-SRB would accelerate the establishment of these microbes in the bioreactors. Eighty-seven days after the start of the flow, the relative abundances of IO-SRB and CO-SRB, respectively, in both inoculated and uninoculated reactors were nearly identical, around 25% and 75%, respectively (Fig. 7A). However, the relative abundances of CO-SRB exceeded those of IO-SRB several weeks earlier in the inoculated reactor (Fig. 7A), after 25 days of pumping, and remained nearly unchanged thereafter. In contrast, CO-SRB became more abundant than the IO-SRB in the uninoculated reactor only after 75 days of pumping (Fig. 7A). The relative abundance of IO-SRB increased with the increased organic loading when mixtures B, C, and D were pumped through the reactors (Fig. 7B). The ratio of IO- to CO-SRB remained relatively constant on consecutively sampled days within each chemical condition, indicating that these shifts reflected a response to the organic load.

**Figure 7:**
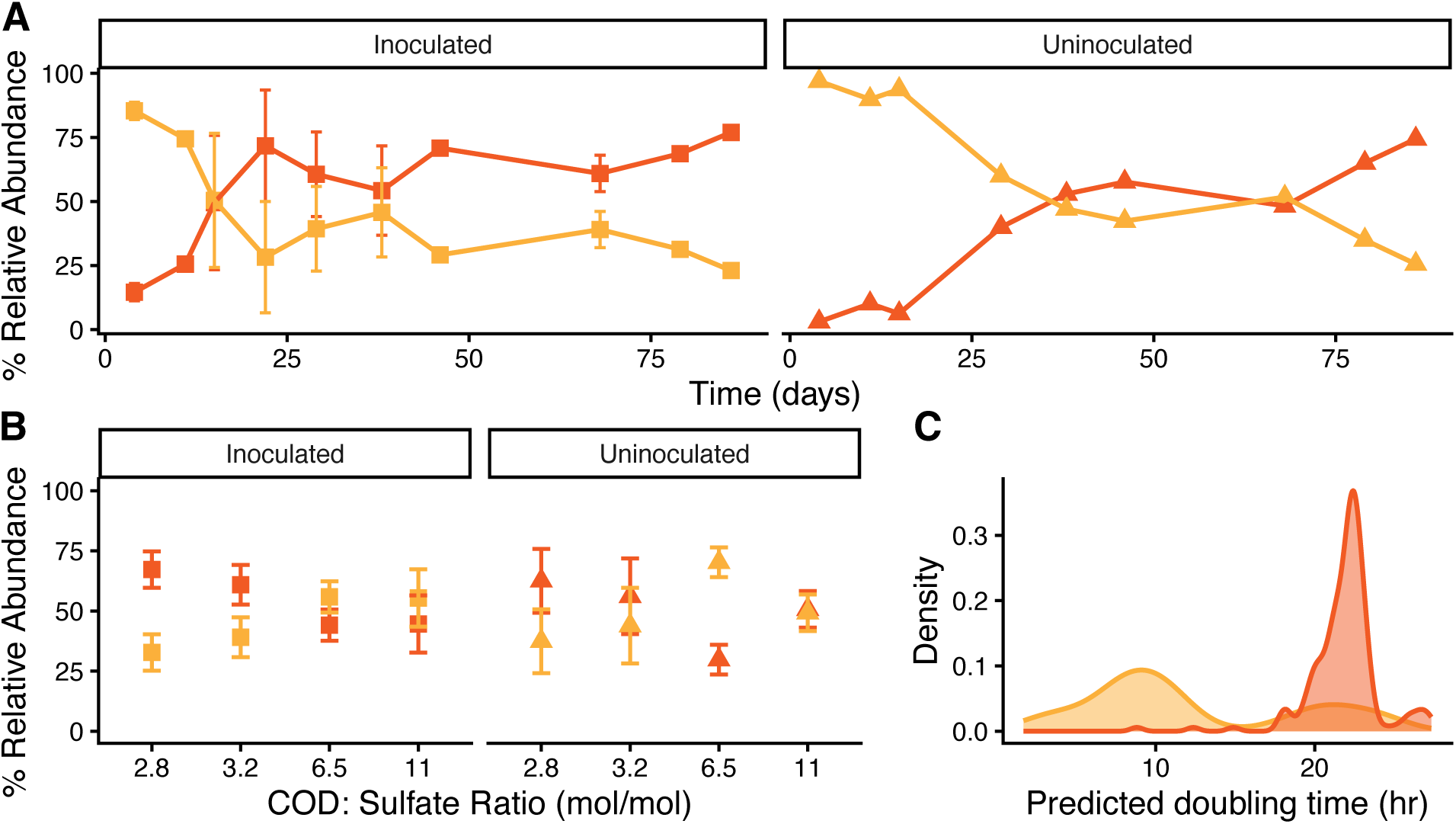
Abundances and predicted doubling times of incomplete- and complete-oxidizing SRB. Across all panels, CO-SRB are indicated in red and IO-SRB in yellow. A) Relative abundance of incomplete-oxidizing (IO-SRB) and complete-oxidizing (CO-SRB) sulfate reducers over time after the onset of pumping, relative to the total SRB population. Error bars denote the standard deviation between duplicate inoculated bioreactors. B) Relative abundance of IO- and CO-SRB under various organic loading ratios (COD: sulfate in mol/mol). Data represent the average compositions from the final three sampling time points for each condition. Values for inoculated reactors are averages of biological duplicates (n ≥ 6), whereas uninoculated values are averages of temporal replicates from the single reactor (n ≥ 3). Error bars show the standard deviation. C) Density distributions of predicted doubling times based on complete genomes of representative IO- and CO-SRB taxa.

## 4. Discussion

To advance the implementation of sulfate-reducing digestion for wastewater treatment, we quantified the kinetics of organic degradation and gypsum conversion in a continuous up-flow anaerobic sludge blanket (UASB) system. This represents a critical step toward scaling up the process to reduce industrial waste and greenhouse gas emissions simultaneously. Here, we benchmark the performance of our system against previously reported flow-through configurations and standard anaerobic digesters and consider factors that control the competition between sulfate-reducing bacteria and methanogens.

### 4.1 Organic removal efficiency in sulfate-containing digesters

Sludge digestion technologies that aim to decrease greenhouse gas emissions must decrease the volume of sludge by oxidizing organic compounds as efficiently as current standards. Organic removal rates in the herein described benchtop sulfate-containing system were 1.5–4.0 g COD/L/day. These rates are comparable to those in traditional sulfate-free anaerobic digestion systems (Musa et al., 2018), including the Deer Island Treatment Plant (Boston, MA) that operates at similar loading rates, but requires heating to mesophilic temperatures (∼37°C) for standard methanogenic digestion.

A further increase in organic removal is likely possible by adjusting thermodynamic parameters. The benchtop system described here operated at room temperature (20–22°C) to reduce the experimental complexity. We anticipate that both the organic removal and the alkalinity production would increase at ∼37°C. This is based on the assumption that the hydrolysis of complex particulate organic matter is the rate-limiting step in sludge digestion that follows the Arrhenius relationship, where enzymatic activity and metabolic rates increase with temperature (van Lier et al., 1997).

### 4.2 Mechanism of methane suppression

Previous work predicted that methane production during the degradation of sewage sludge in the presence of sulfate could be suppressed if complete-oxidizing sulfate-reducing bacteria (CO-SRB) outcompete methanogens for acetate (Coon et al., 2026). Experiments in a continuous flow system confirmed this prediction by achieving high rates of organic and sulfate removal with negligible methane production (∼1% of electron flow) (Fig. 2, Table S1). We attribute this suppression to the competitive exclusion of acetoclastic methanogens by CO-SRB. Acetate accumulated only when sulfate was depleted in sulfate-limited reactors (Fig. 3) or when sulfate reduction was inhibited in batch cultures (Coon et al., 2026). This is consistent with the generally higher acetate affinity of CO-SRB compared to acetoclastic methanogens (Schönheit et al., 1982).

The extent of methane suppression depends on the balance between organic loading and sulfate availability. In organic-rich environments, a continuous surplus of electron donors can relieve competitive pressure, allowing acetoclastic and other types of methanogenesis (Coon et al., 2026). However, when sulfate reducers oxidize organic compounds completely to CO_2_, they prevent the accumulation of acetate as a feedstock for methanogens. This forces methanogens to rely on H_2_/CO_2_, where methanogens are kinetically disadvantaged compared to SRB (Lovley et al., 1982), or on methyl compounds that are less abundant in these environments (Conrad, 2020).

### 4.3 Competition between CO-SRB and IO-SRB

The suppression of methane flux through the competition for acetate depends on the presence and activity of sulfate-reducing bacteria that can oxidize acetate (Coon et al., 2026). The ability to oxidize acetate distinguishes the two groups of SRB. Incomplete-oxidizing SRB (IO-SRB) are only capable of oxidizing substrates into acetate, while complete-oxidizing SRB (CO-SRB) oxidize substrates, including acetate, into CO_2_. We aimed to enrich for CO-SRB that oxidize organics fully to CO_2_ because this pathway oxidizes acetate (Fig. 8). In contrast, incomplete-oxidizing SRB release acetate that enables methanogenesis (Fig. 8). Thus, controlling the abundance and activity of these two SRB groups should determine whether the system suppresses methanogenesis or enables it.

**Figure 8:**
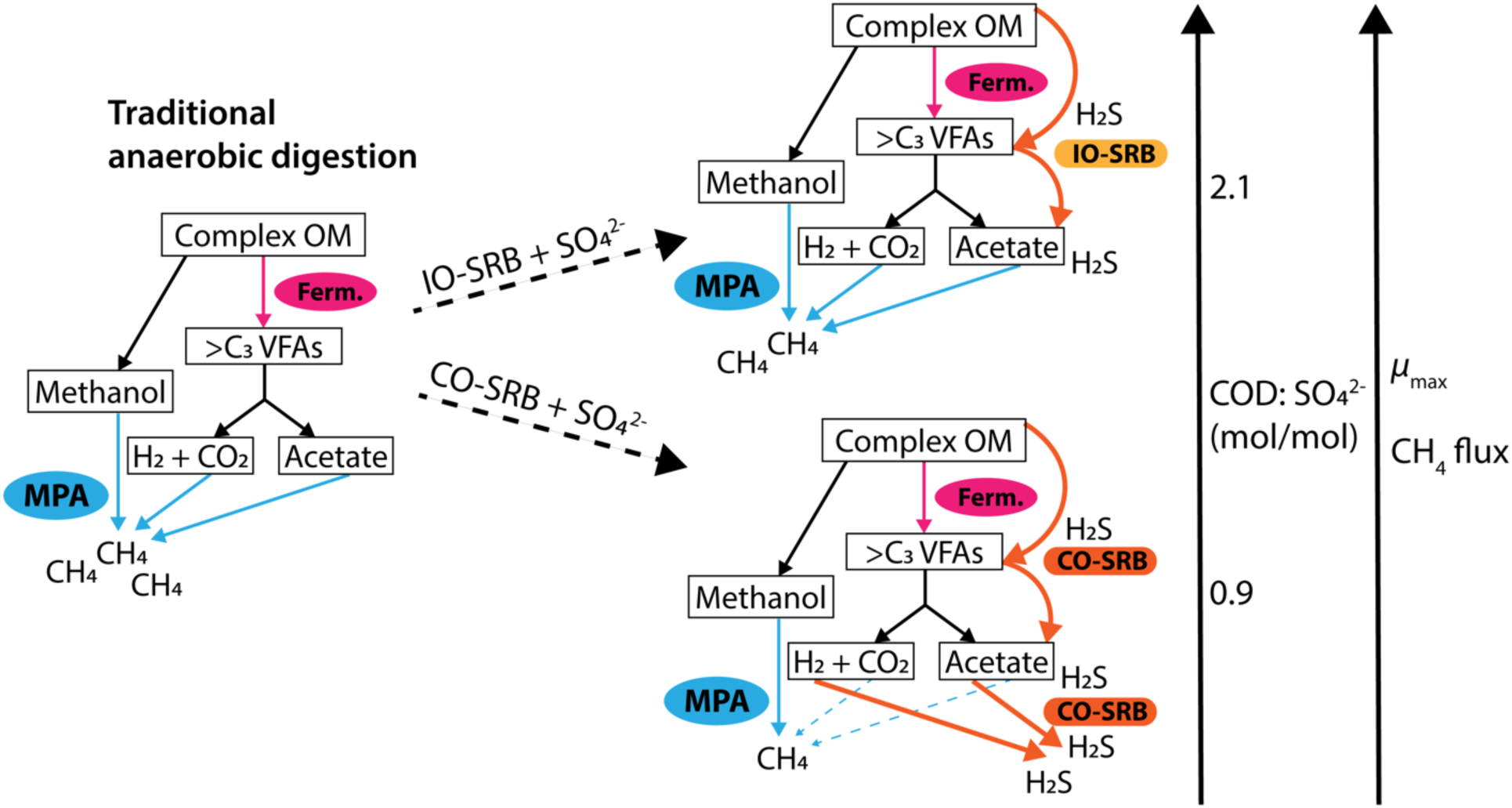
Schematic of metabolic pathways and electron flow in sulfate-reducing digestion. Colored arrows denote specific microbial activities: pink (fermentation), blue (methanogenesis via methane-producing archaea, MPA), yellow (incomplete sulfate reduction via IO-SRB), and orange (complete sulfate reduction via CO-SRB). Abbreviations: OM, organic matter; VFAs, volatile fatty acids; >C3, organic compounds with carbon chains longer than three; Ferm, fermentative microorganisms; μ_max_, maximum specific growth rate.

The influence of organic loading on the ratio of IO- to CO-SRB can be viewed through the lens of the known and predicted growth kinetics of these groups of sulfate-reducing bacteria. At high organic loads, the abundance of various organic substrates favors organisms with faster maximum specific growth rates (*µ*_max_). IO-SRB typically exhibit higher growth rates than CO-SRB when grown on organic compounds such as lactate and ethanol that are produced by the fermentation of more complex organic substrates (Shu et al., 2025; Badziong and Thauer, 1978; Elferink et al., 1998; Kremer et al., 1988; Laanbroek et al., 1983; Nethe-Jaenchen and Thauer, 1984; Traore et al., 1981; Widdel, 1988; McCartney and Oleszkiewicz, 1993; Schönheit et al., 1982). Predicted growth rates of IO-SRB and CO-SRB reflect those experimental constraints (Fig. 7C, from Weissman et al., 2021; Xu et al., 2025).

Consequently, IO-SRB are abundant in nutrient-rich conditions where competition for substrates is less severe. This is also consistent with the stoichiometry of systems that contain substrates such as ethanol, lactate or acetate. Complete oxidation of substrates such as ethanol requires a molar COD: sulfate ratio of ∼0.72, whereas incomplete oxidation occurs at ratios above ∼2.09 (Heijnen and Kleerebezem, 2010; Kleerebezem and Van Loosdrecht, 2010). Thus, limiting or fluctuating sulfate concentrations favor IO-SRB (Cadby et al., 2017; Marietou et al., 2022; Rey et al., 2013). In agreement with these predictions, we observed that the relative abundance of CO-SRB decreased with the increasing COD: sulfate ratio (Fig. 7B), suggesting that high loading rates relaxed the competitive pressure that otherwise favored the more efficient CO-SRB.

The successful enrichment of CO-SRB in a continuous flow system is consistent with recent findings on how reactor setups influence selection (Shu et al., 2025). Shu et al. (2025) observed that pulsed feeding regimes, which establish temporarily high concentrations of substrates, favor the faster-growing IO-SRB. In contrast, continuous flow systems such as the UASB maintain steady, low-substrate concentrations that favor organisms with higher substrate affinities (*µ*_max_/K_s_), such as CO-SRB (Shu et al., 2025). This distinction in operational mode, combined with the experimental duration, also contextualizes results from previous studies. For instance, Dar et al. (2008) observed the dominance of IO-SRB even at a low lactate-to-sulfate ratio of 0.34 mol/mol, a condition that should favor complete oxidation, while noting that their 15-day experiments were likely too short to capture the eventual succession to CO-SRB. By continuing the flow for about 90 days, we confirmed that CO-SRB can overcome the initial kinetic advantage of IO-SRB even in the absence of a starting enrichment that accelerates the establishment and prevalence of CO-SRB in the community. Thus, while high organic loads or pulsed feeding favor IO-SRB, long-term continuous operation selects for CO-SRB. This leads to high rates of organic removal with low methane emissions at molar COD: sulfate ratios below 6.5, with decreased rates of organic removal when the supplied sulfate is fully removed within the reactor at molar COD: sulfate ratios of 11.

### 4.4 Implications for inoculation and community control at scale

Traditional anaerobic digestion transfers active biomass from established digesters to new reactors to accelerate the onset of treatment (Boulanger et al., 2012). This practice requires optimizing the inoculum-to-substrate ratio: higher inoculum volumes reduce the lag time before digestion, while also reducing the reactor volume available for fresh sludge treatment. Our results indicate that similar seeding strategies are applicable to sulfate-reducing digestion. Inoculation accelerated the establishment of a stable community in our experimental system, although the communities in the reactors converged to similar performances and compositions regardless of inoculation (Figs. 2, 4, and S3). The organic loading and sulfate availability were the primary determinants of the composition of communities enriched from the sewage sludge (Fig. 5B).

At Deer Island Wastewater Treatment Plant, the source of sewage sludge used in our experiments, these cycles last ∼24 days, so waiting for ∼90 days for the establishment of a stable community would not be feasible. The addition of a mature enrichment from an active digester should be sufficient to seed a new sulfate-reducing system and accelerate the rates of organic removal and sulfate reduction during each reactor cycle. However, to achieve methane suppression by targeted enrichment of CO-SRB requires biomass retention and management of the initial feed. Future operators should prioritize low COD: sulfate ratios during the start-up phase. This condition, and the addition of inoculum, would accelerate the establishment of slower-growing CO-SRB before the system is exposed to higher organic loads. These higher organic loads should remain below a COD: sulfate ratio of 11 if the hydraulic retention time is ∼12 hours, given that we observed the lowest COD removal rate at these conditions.

## 5. Conclusions

Kinetics of organic degradation and gypsum conversion were quantified in a continuous up-flow anaerobic sludge blanket (UASB) system at room temperature (∼20°C) and a hydraulic retention time of ∼12 hours. The stability of the sulfate-reducing microbial communities in the bioreactors was assessed under continuous flow conditions. The system enabled a ∼70x decrease in electron flow toward methanogenesis compared to standard organic digestion in the absence of sulfate, while maintaining organic removal rates comparable to those of standard municipal systems, such as the Deer Island Treatment Plant (Boston, MA). This process relied on the competitive exclusion of methanogens by SRB that oxidized acetate (CO-SRB). The starting conditions influenced the ratio of complete-oxidizing (CO-SRB) to incomplete-oxidizing (IO-SRB) bacteria. Establishing the sludge blanket at low initial COD: sulfate ratios likely facilitated the enrichment of CO-SRB. Once established, the microbial community remained robust even as organic loads increased. These results enable targeted control of the microbial community and conditions to inform the design of anaerobic digesters to reduce greenhouse gas emissions and remove organic and industrial waste.

## Acknowledgements

This project was supported by the Chan-Zuckerberg and Siegelman funds to the Advanced Carbon Mineralization Initiative (ACMI). GRC acknowledges fellowship funding from GE Vernova. TB acknowledges funding by MIT School of Science Fundamental Science Investigator award. We extend acknowledgement to Lisa Wong and colleagues at Deer Island’s Wastewater Treatment Plant for sample collection. We extend acknowledgement to Maddie Paoletti for help with elemental sulfur measurements.

## Notes

### Competing Interest Statement

The authors have declared no competing interest.

## Works Cited

Alsanea, A., Bounaga, A., Lyamlouli, K., Zeroual, Y., Boulif, R., Zhou, C., Rittmann, B., 2024. Sulfate Leached from Phosphogypsum Is Transformed in a Hydrogen-Based Membrane Biofilm Reactor. ACS EST Eng. 10.1021/acsestengg.4c00563

Badziong, W., Thauer, R.K., 1978. Growth yields and growth rates of Desulfovibrio vulgaris (Marburg) growing on hydrogen plus sulfate and hydrogen plus thiosulfate as the sole energy sources. Arch. Microbiol. 117, 209–214. 10.1007/BF00402310

Bilal, E., Bellefqih, H., Bourgier, V., Mazouz, H., Dumitraş, D.-G., Bard, F., Laborde, M., Caspar, J.P., Guilhot, B., Iatan, E.-L., 2023. Phosphogypsum circular economy considerations: A critical review from more than 65 storage sites worldwide. J. Clean. Prod. 414, 137561.

Boulanger, A., Pinet, E., Bouix, M., Bouchez, T., Mansour, A.A., 2012. Effect of inoculum to substrate ratio (I/S) on municipal solid waste anaerobic degradation kinetics and potential. Waste Manag. 32, 2258–2265. 10.1016/j.wasman.2012.07.024

Bounaga, A., Alsanea, A., Lyamlouli, K., Zhou, C., Zeroual, Y., Boulif, R., Rittmann, B.E., 2022. Microbial transformations by sulfur bacteria can recover value from phosphogypsum: A global problem and a possible solution. Biotechnol. Adv. 57, 107949.

Bueno de Mesquita, C.P., Wu, D., Tringe, S.G., 2023. Methyl-Based Methanogenesis: an Ecological and Genomic Review. Microbiol. Mol. Biol. Rev. 87, e00024–22. 10.1128/mmbr.00024-22

Cadby, I.T., Faulkner, M., Cheneby, J., Long, J., van Helden, J., Dolla, A., Cole, J.A., 2017. Coordinated response of the Desulfovibrio desulfuricans 27774 transcriptome to nitrate, nitrite and nitric oxide. Sci. Rep. 7, 16228. 10.1038/s41598-017-16403-4

Claypool, G.E., Kaplan, I. R., 1974. The Origin and Distribution of Methane in Marine Sediments, in: Kaplan, Isaac R. (Ed.), Natural Gases in Marine Sediments. Springer US, Boston, MA, pp. 99–139. 10.1007/978-1-4684-2757-8_8

Conrad, R., 2020. Importance of hydrogenotrophic, aceticlastic and methylotrophic methanogenesis for methane production in terrestrial, aquatic and other anoxic environments: A mini review. Pedosphere 30, 25–39. 10.1016/S1002-0160(18)60052-9

Coon, G.R., Duesing, P.D., Paul, R., Baily, J.A., Lloyd, K.G., 2023. Biological methane production and accumulation under sulfate-rich conditions at Cape Lookout Bight, NC. Front. Microbiol. 14. 10.3389/fmicb.2023.1268361

Coon, G.R., Kouadio, V., Murphy, C.W.M., Sun, H., Jagoutz, O., Bosak, T., 2026. Organic availability and microbial competition for acetate suppress methane emissions during the conversion of gypsum in sewage sludge. 10.64898/2026.06.20.733556

Coon, G.R., Williams, L.C., Matthews, A., Diaz, R., Kevorkian, R.T., LaRowe, D.E., Steen, A.D., Lapham, L.L., Lloyd, K.G., 2024. Control of hydrogen concentrations by microbial sulfate reduction in two contrasting anoxic coastal sediments. Front. Microbiol. 15. 10.3389/fmicb.2024.1455857

Danouche, M., Bounaga, A., Boulif, R., Zeroual, Y., Benhida, R., Lyamlouli, K., 2023. Optimization of sulfate leaching from Phosphogypsum for efficient bioreduction in a batch bioreactor using a sulfate-reducing microbial consortium. Chem. Eng. J. 475, 146072. 10.1016/j.cej.2023.146072

Dar, S.A., Kleerebezem, R., Stams, A.J.M., Kuenen, J.G., Muyzer, G., 2008. Competition and coexistence of sulfate-reducing bacteria, acetogens and methanogens in a lab-scale anaerobic bioreactor as affected by changing substrate to sulfate ratio. Appl. Microbiol. Biotechnol. 78, 1045–1055. 10.1007/s00253-008-1391-8

de Jong, P., Srinamasivayam, B., Harrison, A., Wardrop, P., Rebsdorf, M., Thorgaard, S., Vale, P., 2026. Methane emissions monitoring at wastewater treatment plants in Europe and Australia. Water Res. X 30, 100480. 10.1016/j.wroa.2025.100480

D’Hondt, S., Rutherford, S., Spivack, A.J., 2002. Metabolic Activity of Subsurface Life in Deep-Sea Sediments. Science 295, 2067–2070. 10.1126/science.1064878

Egger, M., Lenstra, W., Jong, D., Meysman, F.J.R., Sapart, C.J., Veen, C. van der, Röckmann, T., Gonzalez, S., Slomp, C.P., 2016. Rapid Sediment Accumulation Results in High Methane Effluxes from Coastal Sediments. PLOS ONE 11, e0161609. 10.1371/journal.pone.0161609

Elferink, S.J.W.H.O., Luppens, S.B.I., Marcelis, C.L.M., Stams, A.J.M., 1998. Kinetics of Acetate Oxidation by Two Sulfate Reducers Isolated from Anaerobic Granular Sludge. Appl. Environ. Microbiol. 64, 2301–2303. 10.1128/AEM.64.6.2301-2303.1998

Hao, T., Wei, L., Lu, H., Chui, H., Mackey, H.R., van Loosdrecht, M.C.M., Chen, G., 2013. Characterization of sulfate-reducing granular sludge in the SANI® process. Water Res., Microbial ecology of drinking water and wastewater treatment 47, 7042–7052. 10.1016/j.watres.2013.07.052

Heijnen, J.J., Kleerebezem, R., 2010. Bioenergetics of Microbial Growth.

Hu, R., Aronson, H.S., Weaver, M.E., Price, M.N., LaRowe, D.E., Liang, Y., Deutschbauer, A.M., Coates, J.D., Carlson, H.K., 2025. Organic carbon oxidation state shapes fermentative methanogenic microbiomes and controls greenhouse gas fluxes. 10.1101/2025.05.12.653603

Karekar, S., Stefanini, R., Ahring, B., 2022. Homo-Acetogens: Their Metabolism and Competitive Relationship with Hydrogenotrophic Methanogens. Microorganisms 10, 397. 10.3390/microorganisms10020397

Kiene, R.P., Oremland, R.S., Catena, A., Miller, L.G., Capone, D.G., 1986. Metabolism of Reduced Methylated Sulfur Compounds in Anaerobic Sediments and by a Pure Culture of an Estuarine Methanogen. Appl. Environ. Microbiol. 52, 1037–1045. 10.1128/aem.52.5.1037-1045.1986

Kijjanapanich, P., Annachhatre, A.P., Lens, P.N.L., 2014. Biological Sulfate Reduction for Treatment of Gypsum Contaminated Soils, Sediments, and Solid Wastes. Crit. Rev. Environ. Sci. Technol. 44, 1037–1070. 10.1080/10643389.2012.743270

Kleerebezem, R., Van Loosdrecht, M.C.M., 2010. A Generalized Method for Thermodynamic State Analysis of Environmental Systems. Crit. Rev. Environ. Sci. Technol. 40, 1–54. 10.1080/10643380802000974

Kremer, D.R., Nienhuis-Kuiper, H.E., Hansen, T.A., 1988. Ethanol dissimilation in Desulfovibrio. Arch. Microbiol. 150, 552–557. 10.1007/BF00408248

Laanbroek, H.J., Geerligs, H.J., Sijtsma, L., Veldkamp, H., 1983. Competition for Sulfate and Ethanol Among Desulfobacter, Desulfobulbus, and Desulfovibrio Species Isolated from Intertidal Sediments. Appl. Environ. Microbiol. 47, 329–334. 10.1128/aem.47.2.329-334.1984

LaRowe, D., Amend, J., 2014. 13. Energetic constraints on life in marine deep sediments, in: Kallmeyer, J., Wagner, D. (Eds.), Microbial Life of the Deep Biosphere. De Gruyter, pp. 279–302.

LaRowe, D.E., Amend, J.P., 2015. Catabolic rates, population sizes and doubling/replacement times of microorganisms in natural settings. Am. J. Sci. 315, 167–203. 10.2475/03.2015.01

LaRowe, D.E., Arndt, S., Bradley, J.A., Estes, E.R., Hoarfrost, A., Lang, S.Q., Lloyd, K.G., Mahmoudi, N., Orsi, W.D., Shah Walter, S.R., Steen, A.D., Zhao, R., 2020. The fate of organic carbon in marine sediments - New insights from recent data and analysis. Earth-Sci. Rev. 204, 103146. 10.1016/j.earscirev.2020.103146

LaRowe, D.E., Van Cappellen, P., 2011. Degradation of natural organic matter: A thermodynamic analysis. Geochim. Cosmochim. Acta 75, 2030–2042. 10.1016/j.gca.2011.01.020

Lovley, D.R., Dwyer, D.F., Klug, M.J., 1982. Kinetic Analysis of Competition Between Sulfate Reducers and Methanogens for Hydrogen in Sediments. Appl. Environ. Microbiol. 43, 1373–1379. 10.1128/aem.43.6.1373-1379.1982

Lovley, D.R., Klug, M.J., 1983. Methanogenesis from Methanol and Methylamines and Acetogenesis from Hydrogen and Carbon Dioxide in the Sediments of a Eutrophic Lake. Appl. Environ. Microbiol. 45, 1310–1315. 10.1128/aem.45.4.1310-1315.1983

Lu, H., Ekama, G.A., Wu, D., Feng, J., van Loosdrecht, M.C.M., Chen, G.-H., 2012a. SANI® process realizes sustainable saline sewage treatment: Steady state model-based evaluation of the pilot-scale trial of the process. Water Res. 46, 475–490. 10.1016/j.watres.2011.11.031

Lu, H., Wu, D., Jiang, F., Ekama, G.A., van Loosdrecht, M.C.M., Chen, G.-H., 2012b. The demonstration of a novel sulfur cycle-based wastewater treatment process: Sulfate reduction, autotrophic denitrification, and nitrification integrated (SANI®) biological nitrogen removal process. Biotechnol. Bioeng. 109, 2778–2789. 10.1002/bit.24540

Marietou, A., Kjeldsen, K.U., Glombitza, C., Jørgensen, B.B., 2022. Response to substrate limitation by a marine sulfate-reducing bacterium. ISME J. 16, 200–210. 10.1038/s41396-021-01061-2

McCartney, D.M., Oleszkiewicz, J.A., 1993. Competition between methanogens and sulfate reducers: effect of COD:sulfate ratio and acclimation. Water Environ. Res. 65, 655–664. 10.2175/WER.65.5.8

Murphy, C.W.M., Coon, G.R., Jagoutz, O., Bosak, T., 2026. Enrichment of sulfate-reducing microbial communities for sewage sludge treatment. Chem. Eng. J. Green Sustain. 100054. 10.1016/j.cejgas.2026.100054

Musa, M.A., Idrus, S., Hasfalina, C.M., Daud, N.N.N., 2018. Effect of Organic Loading Rate on Anaerobic Digestion Performance of Mesophilic (UASB) Reactor Using Cattle Slaughterhouse Wastewater as Substrate. Int. J. Environ. Res. Public. Health 15, 2220. 10.3390/ijerph15102220

Nethe-Jaenchen, R., Thauer, R.K., 1984. Growth yields and saturation constant of Desulfovibrio vulgaris in chemostat culture. Arch. Microbiol. 137, 236–240. 10.1007/BF00414550

Omil, F., Lens, P., Hulshoff Pol, L., Lettinga, G., 1996. Effect of upward velocity and sulphide concentration on volatile fatty acid degradation in a sulphidogenic granular sludge reactor. Process Biochem. 31, 699–710. 10.1016/S0032-9592(96)00015-5

Oremland, R.S., Taylor, B.F., 1978. Sulfate reduction and methanogenesis in marine sediments. Geochim. Cosmochim. Acta 42, 209–214. 10.1016/0016-7037(78)90133-3

Oshita, K., Okumura, T., Takaoka, M., Fujimori, T., Appels, L., Dewil, R., 2014. Methane and nitrous oxide emissions following anaerobic digestion of sludge in Japanese sewage treatment facilities. Bioresour. Technol. 171, 175–181. 10.1016/j.biortech.2014.08.081

Ozuolmez, D., Na, H., Lever, M.A., Kjeldsen, K.U., Jørgensen, B.B., Plugge, C.M., 2015. Methanogenic archaea and sulfate reducing bacteria co-cultured on acetate: teamwork or coexistence? Front. Microbiol. 6. 10.3389/fmicb.2015.00492

Plyatsuk, L., Chernish, E., 2014. Intensification of Anaerobic Microbiological Degradation of Sewage Sludge and Gypsum Waste Under Bio-Sulfidogenic Conditions. J. Solid Waste Technol. Manag. 40, 10–23. 10.5276/JSWTM.2014.10

Plyatsuk, L.D., Chernysh, Y.Y., 2016. The Removal of Hydrogen Sulfide in the Biodesulfurization System Using Granulated Phosphogypsum. Eurasian Chem.-Technol. J. 18, 47–54. 10.18321/ectj395

Reeburgh, W.S., 2007. Oceanic Methane Biogeochemistry. Chem. Rev. 107, 486–513. 10.1021/cr050362v

Rey, F.E., Gonzalez, M.D., Cheng, J., Wu, M., Ahern, P.P., Gordon, J.I., 2013. Metabolic niche of a prominent sulfate-reducing human gut bacterium. Proc. Natl. Acad. Sci. U. S. A. 110, 13582–13587. 10.1073/pnas.1312524110

Rzeczycka, M., Miernik, A., Markiewicz, Z., 2010. Simultaneous Degradation of Waste Phosphogypsum and Liquid Manure from Industrial Pig Farm by a Mixed Community of Sulfate-Reducing Bacteria. Pol. J. Microbiol. 59, 241–247. 10.33073/pjm-2010-037

Salo, M., Mäkinen, J., Yang, J., Kurhila, M., Koukkari, P., 2018. Continuous biological sulfate reduction from phosphogypsum waste leachate. Hydrometallurgy 180, 1–6. 10.1016/j.hydromet.2018.06.020

Schönheit, P., Kristjansson, J.K., Thauer, R.K., 1982. Kinetic mechanism for the ability of sulfate reducers to out-compete methanogens for acetate. Arch. Microbiol. 132, 285–288. 10.1007/BF00407967

Sela-Adler, M., Ronen, Z., Herut, B., Antler, G., Vigderovich, H., Eckert, W., Sivan, O., 2017. Co-existence of Methanogenesis and Sulfate Reduction with Common Substrates in Sulfate-Rich Estuarine Sediments. Front. Microbiol. 8. 10.3389/fmicb.2017.00766

Shu, W., Du, B., Wu, G., 2025. Strategies for enriching targeted sulfate-reducing bacteria and revealing their microbial interactions in anaerobic digestion ecosystems. Water Res. 270, 122842.

Traore, A.S., Hatchikian, C.E., Gall, J.L., Belaich, J.-P., 1981. Microcalorimetric studies of the growth of sulfate-reducing bacteria: comparison of the growth parameters of some Desulfovibrio species. J. Bacteriol. 149, 606–611. 10.1128/jb.149.2.606-611.1982

Uberoi, V., Bhattacharya, S.K., 1997. Sulfate-Reducing Bacteria in Anaerobic Propionate Systems. J. Environ. Eng. 123, 675–682. 10.1061/(ASCE)0733-9372(1997)123:7(675)

van Lier, J.B., Rebac, S., Lettinga, G., 1997. High-rate anaerobic wastewater treatment under psychrophilic and thermophilic conditions. Water Sci. Technol. 35, 199–206. 10.2166/wst.1997.0383

Wang, J., Lu, H., Chen, G.-H., Lau, G.N., Tsang, W.L., van Loosdrecht, M.C.M., 2009. A novel sulfate reduction, autotrophic denitrification, nitrification integrated (SANI) process for saline wastewater treatment. Water Res. 43, 2363–2372. 10.1016/j.watres.2009.02.037

Wang, J., Shi, M., Lu, H., Wu, D., Shao, M.-F., Zhang, T., Ekama, G.A., van Loosdrecht, M.C.M., Chen, G.-H., 2011. Microbial community of sulfate-reducing up-flow sludge bed in the SANI® process for saline sewage treatment. Appl. Microbiol. Biotechnol. 90, 2015–2025. 10.1007/s00253-011-3217-3

Weissman, J.L., Hou, S., Fuhrman, J.A., 2021. Estimating maximal microbial growth rates from cultures, metagenomes, and single cells via codon usage patterns. Proc. Natl. Acad. Sci. 118, e2016810118. 10.1073/pnas.2016810118

Widdel, F., 1988. Microbiology and ecology of sulfate-and sulfur-reducing bacteria.

Wu, D., Ekama, G.A., Chui, H.-K., Wang, B., Cui, Y.-X., Hao, T.-W., van Loosdrecht, M.C.M., Chen, G.-H., 2016. Large-scale demonstration of the sulfate reduction autotrophic denitrification nitrification integrated (SANI®) process in saline sewage treatment. Water Res. 100, 496–507. 10.1016/j.watres.2016.05.052

Xiao, K.-Q., Beulig, F., Kjeldsen, K.U., Jørgensen, B.B., Risgaard-Petersen, N., 2017. Concurrent Methane Production and Oxidation in Surface Sediment from Aarhus Bay, Denmark. Front. Microbiol. 8. 10.3389/fmicb.2017.01198

Xiao, K.-Q., Beulig, F., Røy, H., Jørgensen, B.B., Risgaard-Petersen, N., 2018. Methylotrophic methanogenesis fuels cryptic methane cycling in marine surface sediment. Limnol. Oceanogr. 63, 1519–1527. 10.1002/lno.10788

Xu, L., Zakem, E., Weissman, J.L., 2025. Improved maximum growth rate prediction from microbial genomes by integrating phylogenetic information. Nat. Commun. 16, 4256. 10.1038/s41467-025-59558-9

Yamamoto, S., Alcauskas, J.B., Crozier, T.E., 1976. Solubility of methane in distilled water and seawater. J. Chem. Eng. Data 21, 78–80. 10.1021/je60068a029

Zhang, M., Wang, H., 2014. Organic wastes as carbon sources to promote sulfate reducing bacterial activity for biological remediation of acid mine drainage. Miner. Eng. 69, 81–90. 10.1016/j.mineng.2014.07.010

Zhang, Zhao, Zhang, C., Yang, Y., Zhang, Zhuowei, Tang, Y., Su, P., Lin, Z., 2022. A review of sulfate-reducing bacteria: Metabolism, influencing factors and application in wastewater treatment. J. Clean. Prod. 376, 134109. 10.1016/j.jclepro.2022.134109

Zhuang, G.-C., Elling, F.J., Nigro, L.M., Samarkin, V., Joye, S.B., Teske, A., Hinrichs, K.-U., 2016. Multiple evidence for methylotrophic methanogenesis as the dominant methanogenic pathway in hypersaline sediments from the Orca Basin, Gulf of Mexico. Geochim. Cosmochim. Acta 187, 1–20. 10.1016/j.gca.2016.05.005

Zhuang, G.-C., Heuer, V.B., Lazar, C.S., Goldhammer, T., Wendt, J., Samarkin, V.A., Elvert, M., Teske, A.P., Joye, S.B., Hinrichs, K.-U., 2018. Relative importance of methylotrophic methanogenesis in sediments of the Western Mediterranean Sea. Geochim. Cosmochim. Acta 224, 171–186. 10.1016/j.gca.2017.12.024

