## Supplemental Material for "Minimizing methane emissions during the degradation of sewage sludge in a sulfate-rich bioreactor"

**Supplemental materials**

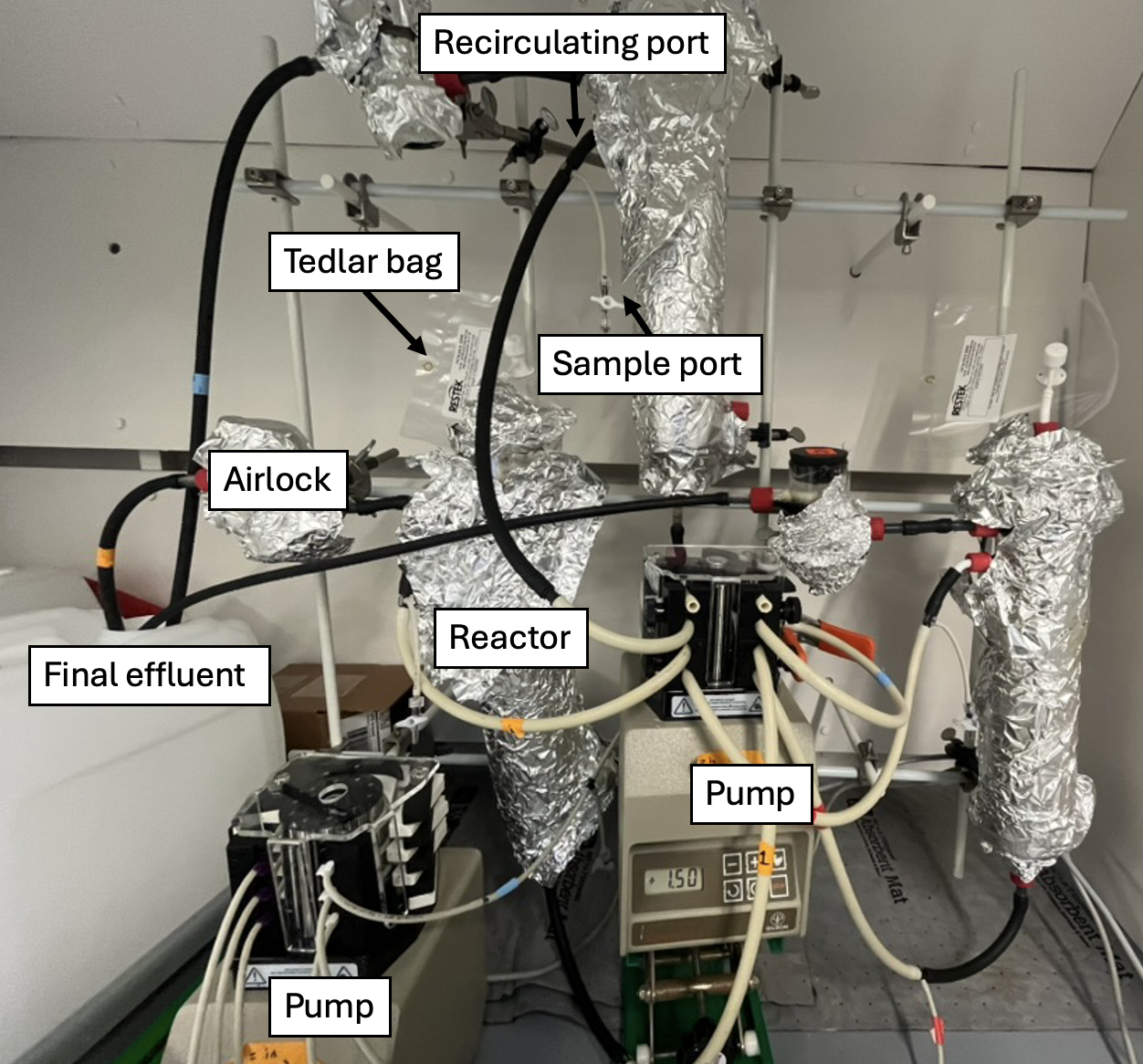

*Figure S1: Reactor while covered by aluminum foil.*

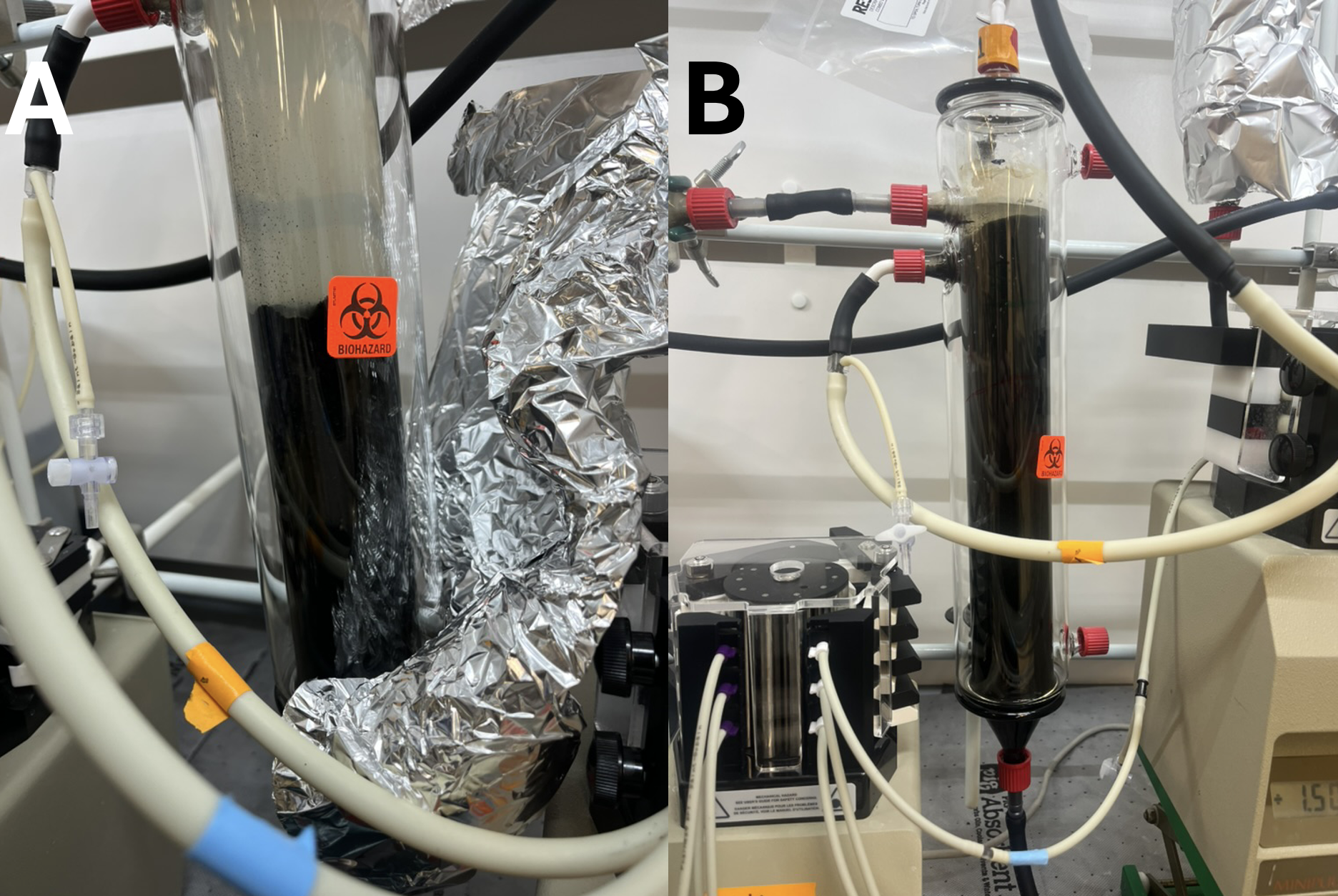

*Figure S2: Sludge blanket formation in BR1. A) day 11 of continuous operation, B) day 37.*

*Table S1: Summary of conditions and important performance metrics.*

| COD: SO_4_^2-^ (mol/mol) | Inoculated? | COD removal | COD removal rate (g/L/d) | Sulfate reduced | Sulfate reduction rate (g/L/d) | Methane production rate (g/L/d) |
| --- | --- | --- | --- | --- | --- | --- |
| 3.24 (n=10) | Yes | 66% | 2.33 | 46% | 1.37 | 0.009 |
| 3.24 (n=5) | No | 86% | 2.67 | 34% | 0.89 | 0.006 |
| 6.45 (n=6) | Yes | 51% | 3.75 | 86% | 1.59 | 0.006 |
| 6.45 (n=3) | No | 63% | 4.06 | 83% | 1.44 | 0.006 |
| 11.0 (n=4) | Yes | 19% | 1.60 | 87% | 0.96 | 0.013 |
| 11.0 (n=2) | No | 21% | 1.75 | 89% | 1.05 | 0.006 |

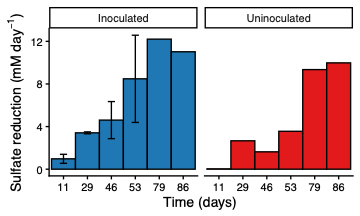

*Figure S3: Sulfate reduction rates in inoculated and uninoculated reactors at COD: sulfate 2.8 mol/mol during the accumulation of sludge blanket and establishment of a stable sludge blanket. Error bars represent duplicate reactors. Time marks days after the initial onset of pumping.*

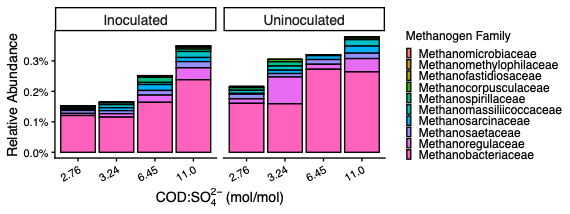

*Figure S4: Methanogenic abundances based on 16S rRNA gene sequencing. Data represent the average compositions at the final three sampling times for each condition*
